# Lytic transglycosylase repertoire diversity enables daughter cell separation and antibiotic resistance in *Escherichia coli* under acidic stress

**DOI:** 10.1101/2024.02.16.580723

**Authors:** Ji Eun Son, Si Hyoung Park, Umji Choi, Chang-Ro Lee

## Abstract

Peptidoglycan (PG) is an indispensable architectural element that imparts physical toughness and rigidity to the bacterial envelope. It is also a dynamic structure that undergoes continuous turnover or autolysis. *Escherichia coli* possesses redundant autolytic enzymes responsible for PG turnover; however, the rationale behind the existence of numerous autolytic enzymes remains incompletely understood. In this study, we elucidated the physiological roles of MltE and MltC, members of the lytic transglycosylase (LTG) family that catalyze the cleavage of glycosidic bonds between disaccharide subunits within PG strands. MltE and MltC are acidic LTGs that exhibit increased enzymatic activity and protein levels under acidic pH conditions, respectively. Deletion of these two LTGs results in a pronounced growth defect and elevated membrane permeability at acidic pH. Furthermore, these two LTGs are crucial for resistance against various antibiotics, particularly vancomycin. Intriguingly, inactivation of these LTGs induces a chaining morphology, indicative of daughter cell separation defects, only under acidic pH conditions. Simultaneous deletion of PG amidases, the known contributors to daughter cell separation, exacerbates the chaining phenotype at acidic pH. This suggests that the two LTGs may participate in the cleavage of glycan strands between daughter cells that cannot be resolved by PG amidases under acidic pH conditions. Collectively, our findings highlight the role of LTG repertoire diversity in facilitating bacterial survival and antibiotic resistance under stressful conditions.

**IMPORTANCE:** Orchestration of peptidoglycan (PG) synthesis and degradation plays a critical role in bacterial growth and development of antibiotic resistance. Bacteria use a diverse array of enzymes involved in PG synthesis and degradation to navigate the challenging conditions of the periplasm. Although *Escherichia coli* harbors 12 lytic transglycosylases (LTGs) responsible for cleaving glycosidic bonds between disaccharide subunits within PG strands, the physiological significance of their redundancy remains unclear. In this study, we demonstrated the indispensability of two LTGs, MltE and MltC, for cell growth, daughter cell separation, and antibiotic resistance under acidic stress conditions. Our findings highlight the potential significance of MltE and MltC as primary targets for the development of potentiators to augment the antimicrobial efficacy of antibiotics in acidic pH environments.

**One sentence summary:** Acid stress-specific lytic transglycosylases

## INTRODUCTION

Bacteria frequently encounter diverse harsh environmental stress conditions that can disrupt pivotal enzymatic reactions, leading to the cessation of bacterial growth. In response to these challenges, bacteria employ a variety of precise survival strategies, and the integrity of the cytoplasmic membrane is a critical component of this defense mechanism. The cytoplasmic membrane serves a crucial role as a barrier preventing the intrusion of toxic molecules into the cytosol. As a result, physiological conditions within the cytoplasm contribute to the maintenance of homeostasis, even in the face of adverse extracellular environments. In Gram-negative bacteria, the periplasm represents a unique space where the permeation of low-molecular-weight toxic molecules occurs readily through the outer membrane porins. However, despite this vulnerability, the periplasm is also the site where essential enzymatic reactions, such as peptidoglycan synthesis, must occur. Consequently, Gram-negative bacteria must devise specialized strategies to ensure the success of periplasmic enzymatic reactions under challenging conditions imposed by harsh environmental stress.

Peptidoglycan (PG) is a mesh-like structure that imparts physical toughness and rigidity to bacteria. The two final enzymatic reactions in its biosynthesis pathway, namely transglycosylation and transpeptidation, occur within the periplasm. Consequently, proteins possessing these enzymatic activities must employ strategies to navigate stress-labile conditions of the periplasm. A recent study illuminated the role of class A penicillin-binding proteins (PBP) 1a and 1b in overcoming specific environmental challenges, with PBP1a facilitating resilience under alkaline conditions, and PBP1b enabling the adaptation to acidic environments (1).

Bacteria possess numerous PG hydrolases and autolysins that function in the periplasm. One of these enzymes is lytic transglycosylase (LTG), which cleaves the β-1,4-glycosidic linkages between N-acetylglucosamine and N-acetylmuramic acid within PG strands. LTGs are involved in various physiological processes, including the termination of PG glycan polymerization (2–4), degradation of synthesis-derived peptidoglycan polymers (5), daughter cell separation (6, 7), PG quality control (8), and insertion of PG-spanning protein complexes, such as flagellar and type VI secretion systems (9, 10). *Escherichia coli* harbors 12 enzymes with LTG activity, indicating a high redundancy of LTGs. Other PG-cleaving enzymes in the periplasm also exhibit redundancy, with some specialized in PG cleavage under periplasmic stress conditions (11–14). Although LTGs display the highest redundancy rate among the PG-cleaving enzymes, studies providing a fundamental understanding of the underlying mechanisms are scarce.

In this study, we elucidated the physiological role of the LTG repertoire diversity. We demonstrated that two specific LTGs, MltE and MltC, are essential for the growth and daughter cell separation of *E. coli* under acidic pH conditions. Both MltE and MltC are categorized as acidic LTGs, as their enzymatic activity and protein stability are significantly augmented in response to acidic stress, respectively. The mutant strain lacking both LTGs exhibited enhanced susceptibility to several antibiotics, including vancomycin, only under acidic pH conditions. Taken together, our data highlight how the diversity of LTGs enables *E. coli* to grow, perform daughter cell separation, and exhibit antibiotic resistance under acidic stress.

## RESULTS

### MltE is required for cell growth at acidic pH

To identify the LTGs crucial for cell growth under acidic pH conditions, we analyzed the cell growth of 12 LTG-defective mutant strains at low pH. While most mutant strains showed growth comparable to that of the wild-type strain in LB medium at pH 4.7, the *mltE* mutant displayed a significant growth defect (Fig. 1A). This phenotype was restored by ectopic expression of MltE (Fig. 1B), indicating that the growth defect was caused by the deletion of the *mltE* gene. Glutamine substitution of the active site glutamate residue at position 64 results in the loss of the LTG activity of MltE (15). Ectopic expression of MltE(E64Q) did not complement the phenotype of the *mltE* mutant (Fig. 1B), suggesting that the LTG activity of MltE is required for cell growth at acidic pH. We then determined if its localization to the outer membrane (16) is important for its cellular function, by constructing two mutant proteins: DsbA(ss)-MltE with the signal sequence of the periplasmic protein DsbA at the N-terminus of MltE to localize it to the periplasm (17) and MltE(S17D&S18D), whose outer membrane-targeted signal (Ser^2+^-Ser^3+^) was replaced with the inner membrane retention signal (Asp^2+^-Asp^3+^) (18). The acid sensitivity of the *mltE* mutant was fully complemented by the ectopic expression of wild-type MltE but not by DsbA(ss)-MltE or MltE(S17D&S18D) (Fig. 1C), indicating that the outer membrane localization of MltE is important for its cellular function. Collectively, these results demonstrate that the LTG activity of MltE in the outer membrane is indispensable for bacterial growth under acidic pH conditions.

**FIG 1.**
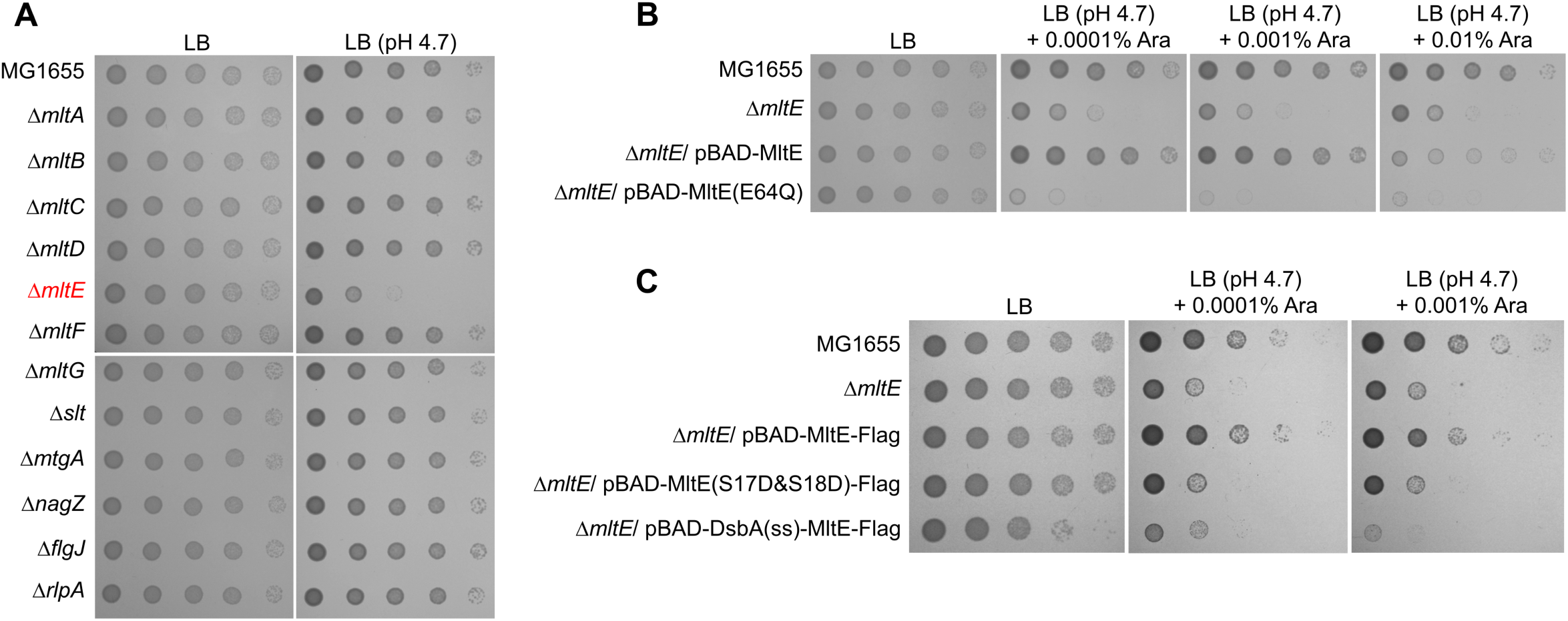
The enzymatic activity of MltE is required for bacterial growth at acidic pH. (A) Growth defect of the *mltE* mutant at acidic pH. The wild-type and indicated mutant cells were serially diluted from 10^8^ to 10^4^ cells/ml in 10-fold steps and spotted onto LB plates or LB plates of pH 4.7. (B and C) Cells of the indicated strains were serially diluted from 10^8^ to 10^4^ cells/ml in 10-fold steps and spotted onto an LB plate and acidified LB plates at pH 4.7 containing the indicated concentrations of arabinose (Ara). (B) Requirement of the enzymatic activity of MltE for the cell growth at acidic pH. (C) The importance of MltE localization to its function.

### The LTG activity of MltE increases at acidic pH

Given that the enzymatic activity of MltE is crucial for cellular growth under acidic pH conditions, we investigated whether its expression and enzymatic activity increase in an acidic environment. First, we examined the mRNA levels of the *mltE* gene at acidic and neutral pH and found them to be comparable (Fig. 2A). Next, we measured the protein level of MltE and found it to be unchanged at acidic pH compared to neutral pH (Fig. 2B). Similar to MltE, the protein level of another LTG MltG at acidic pH was comparable to that at neutral pH (Fig. 2C). Finally, we examined the enzymatic activity of MltE at various pH values. The LTG activity of MltE was measured using purified MltE and PG sacculi covalently labeled with the Remazol brilliant blue (RBB) dye, as previously reported (2, 19). The amount of RBB dye released into the supernatant upon PG cleavage was determined by measuring the absorbance at 595 nm. Notably, the amount of RBB dye released by MltE increased at pH 5.5 and 5.0 compared with that at pH 7.0 (Fig. 2D). RBB dye release was almost undetectable in the presence of MltE(E64Q), indicating that RBB dye release was caused by the LTG activity of MltE. Unlike MltE, the enzymatic activity of MltG diminished at acidic pH compared to that at pH 7.0 (Fig. 2E). Collectively, these results demonstrate that the LTG activity of MltE increases at acidic pH.

**FIG 2.**
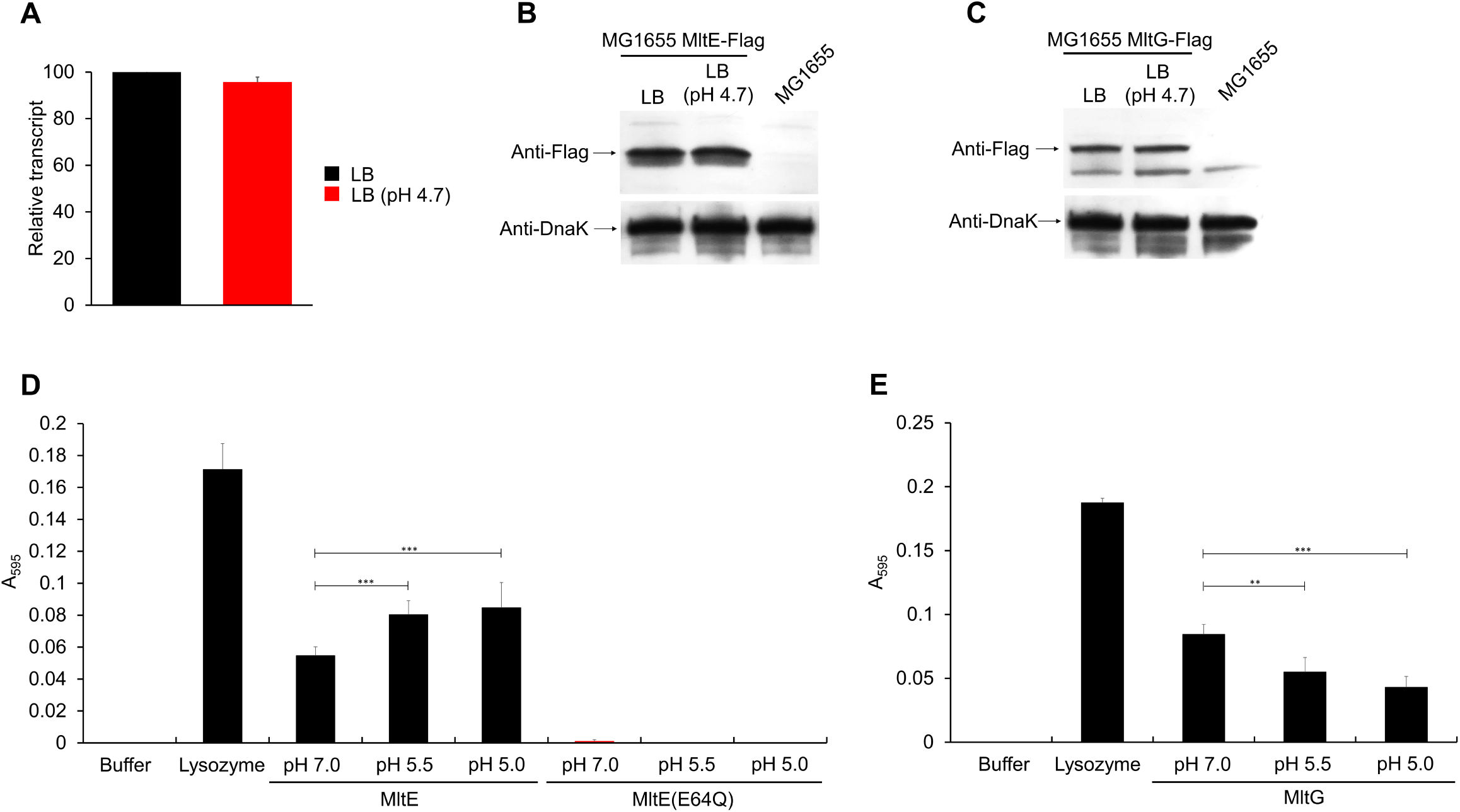
Enhanced enzymatic activity of MltE at acidic pH. (A) Relative mRNA levels of the *mltE* gene in LB medium (black bar) and LB medium of pH 4.7 (red bar). Total mRNA was extracted from the wild-type cells grown to the early exponential phase (OD_600_ = 0.4) in the indicated media. mRNA levels of the *mltE* gene were normalized to the levels of 16S rRNA. Data were produced from three independent experiments. (B and C) Protein levels of MltE (B) and MltG (C) in LB medium or acidified LB medium at pH 4.7. The protein levels were measured using MG1655 MltE-Flag and MltG-Flag cells grown to the early exponential phase (OD_600_ = 0.4) in the indicated media. DnaK was used as the loading control. The exact positions of MltE and MltG were determined by Western blot analysis (MG1655) with the anti-Flag antibody in MG1655 cells as a control. (D and E) Assessment of enzymatic activities of MltE (D) and MltG (E) at acidic pHs. The LTG activity was measured using RBB-labeled PG and purified proteins. Purified protein (500 μg/ml) was incubated at 37°C for 120 min in 50 mM potassium phosphate (pH 7.0) or 50 mM potassium citrate (pH 5.5 or 5.0) buffer containing RBB-labeled PG. The amount of RBB released by LTGs was measured at 595 nm. Lysozyme and MltE(E64Q) were used as the negative and positive control, respectively. Data were obtained from three independent experiments. \*\**p*<0.01; \*\*\**p*<0.001.

### MltE confers vancomycin resistance at acidic pH

To investigate the additional roles of MltE under acidic stress, we assessed its impact on antibiotic resistance at acidic pH. In Mueller-Hinton (MH) broth, the depletion of MltE did not influence the Minimum Inhibitory Concentrations (MICs) of all tested antibiotics (Fig. 3A). However, in MH broth at pH 5.2, the *mltE* mutant exhibited different susceptibilities to various antibiotics. Particularly noteworthy was the 8-fold reduction in the MIC of vancomycin in the *mltE* mutant compared with that in the wild-type strain (Fig. 3A). The sensitivity of the *mltE* mutant to vancomycin at acidic pH was restored by the ectopic expression of MltE (Fig. 3B). These results suggest that MltE is essential for vancomycin resistance under acidic pH conditions.

**FIG 3.**
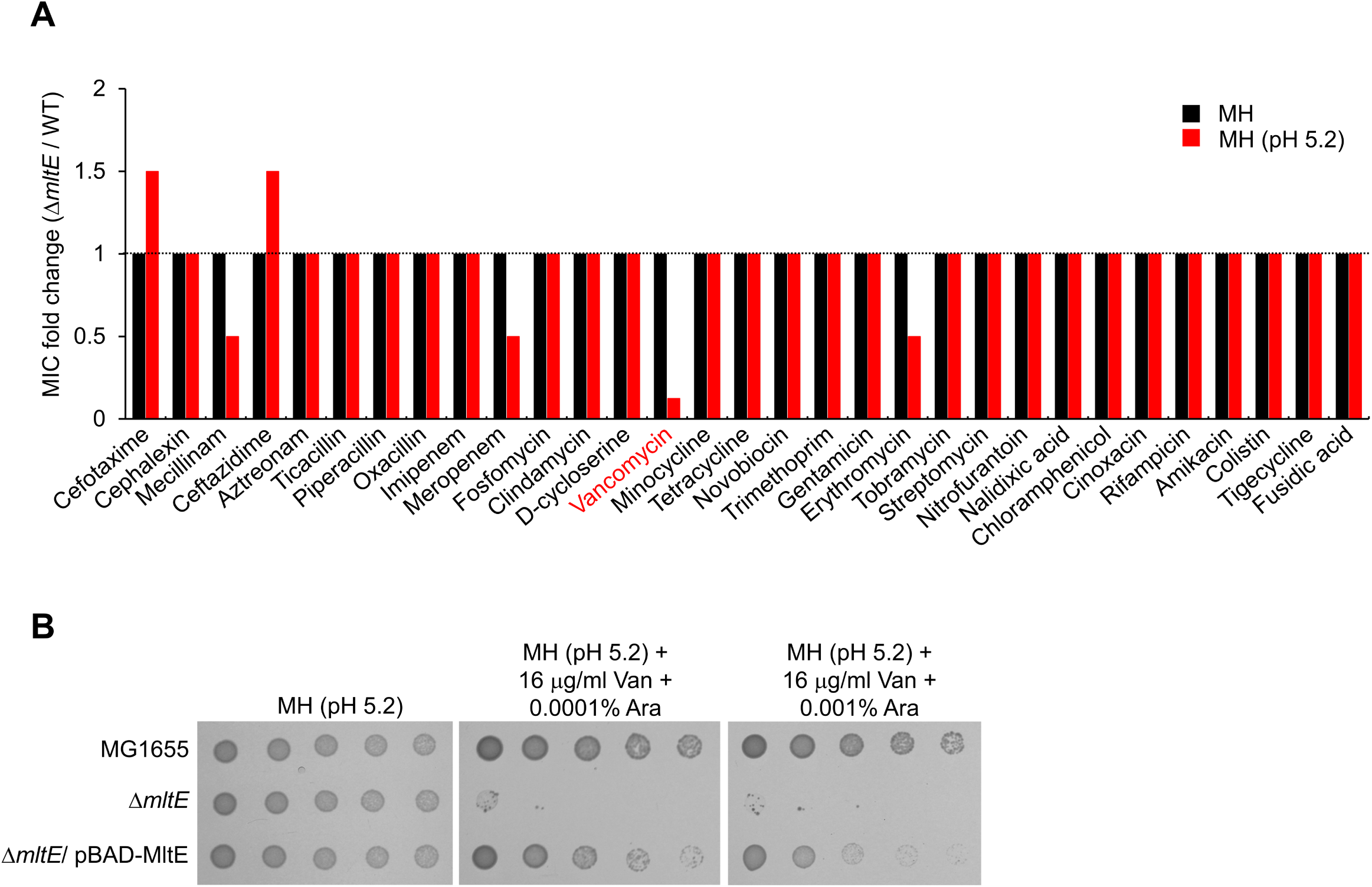
Effect of MltE on antibiotic resistance at acidic pH. (A) Effect of MltE on the MICs of antibiotics. The MICs of various antibiotics were measured against the wild-type and *mltE* mutant strains in MH medium (black bars) or acidified MH medium at pH 5.2 (red bars). The relative MIC values for the *mltE* mutant cells compared to those for the wild-type cells are presented. (B) Complementation of vancomycin sensitivity of the *mltE* mutant. The cells of the indicated strains were serially diluted from 10^8^ to 10^4^ cells/ml in 10-fold steps and spotted onto an MH plate at pH 5.2 and MH plates at pH 5.2 containing 16 μg/ml vancomycin and the indicated concentrations of arabinose (Ara).

### The protein level of MltC increases at acidic pH

The presence of 12 LTGs in *E. coli* prompted us to investigate whether there are additional LTGs that function at acidic pH. To identify other LTGs associated with overcoming acidic stress, we examined which LTGs could complement the phenotype of the *mltE* mutant through ectopic expression. In addition to MltE, only MltC partially complemented the acid sensitivity of the *mltE* mutant (Fig. 4A), indicating that MltC may also regulate cell growth under acidic pH conditions. Notably, the protein level of MltC increased more than two-fold in LB medium at acidic pH compared to that in LB medium at neutral pH (Figs. 4B and C). These results imply that MltC plays a physiological role under acidic conditions.

**FIG 4.**
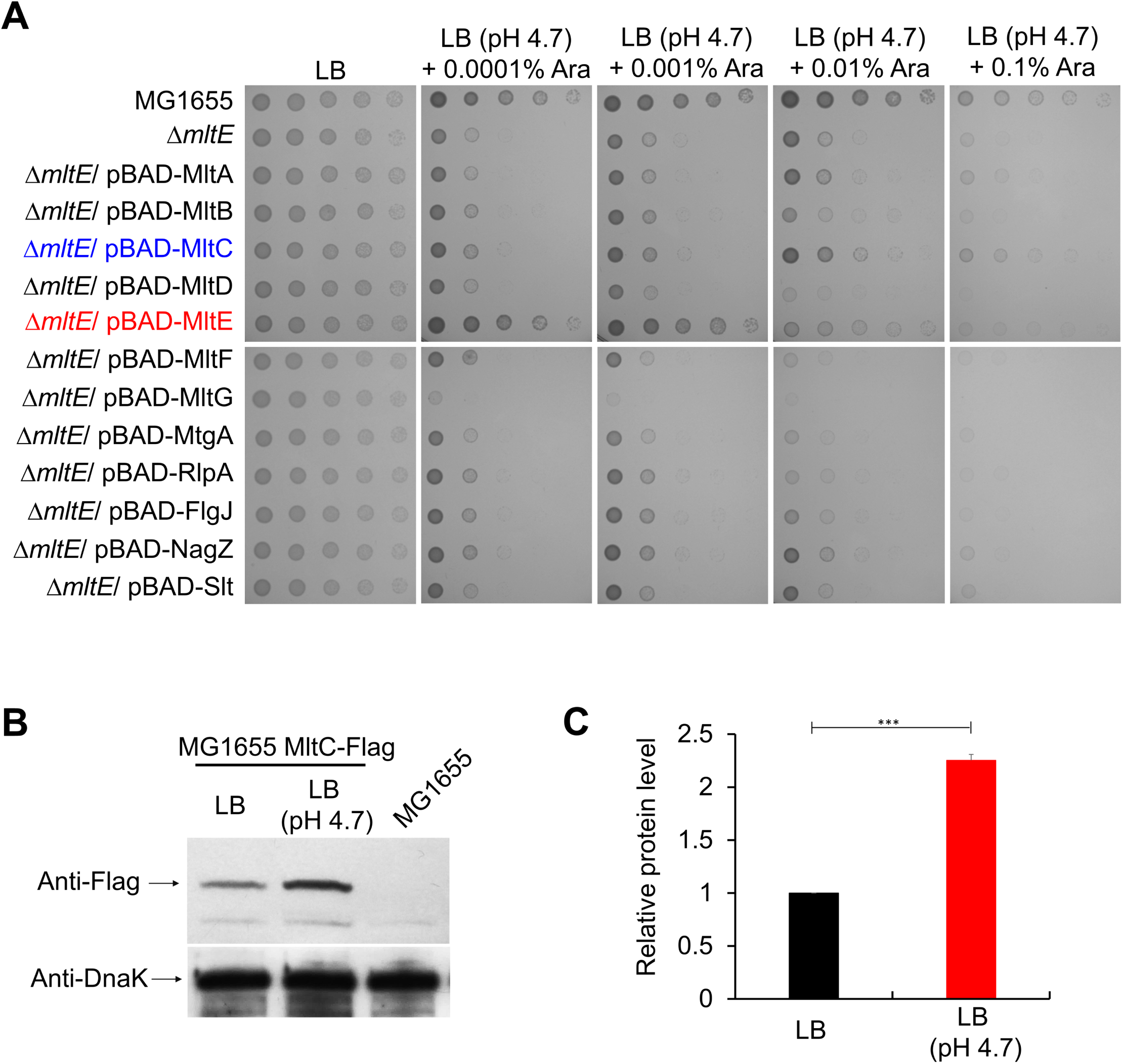
Increased protein level of MltC at acidic pH. (A) Complementation of the growth defect of the *mltE* mutant at acidic pH. Cells of the indicated strains were serially diluted from 10^8^ to 10^4^ cells/ml in 10-fold steps and spotted onto an LB plate and acidified LB plates at pH 4.7 containing the indicated concentrations of arabinose (Ara). (B) Protein levels of MltC in LB medium or acidified LB medium at pH 4.7. The protein levels were measured using MG1655 MltC-Flag cells grown to the early exponential phase (OD_600_ = 0.4) in the indicated media. DnaK was used as the loading control. The exact positions of MltE and MltG were determined by Western blot analysis (MG1655) with the anti-Flag antibody in MG1655 cells as a control. (C) Quantification of the protein levels in (B) plotted as relative levels: black bar, LB; red bar, LB at pH 4.7. Error bars represent the standard deviation from triplicate measurements. \*\*\**p*<0.001.

### MltE and MltC are indispensable for bacterial growth and antibiotic resistance at acidic pH

To analyze the role of MltC at acidic pH, we generated an *mltE mltC* double mutant. Although depletion of MltC alone did not affect bacterial growth at acidic pH (Fig. 1A), additional depletion of MltC in the *mltE* mutant significantly exacerbated its growth defect at acidic pH (Fig. 5A). The *mltE* mutant displayed a severe growth defect at pH 4.7, but showed nearly normal growth comparable to that of the wild-type strain at pH 5.2. In contrast, the *mltE mltC* double mutant exhibited a severe growth defect, even at pH 5.2. The growth defect of the *mltE mltC* double mutant at pH 5.2 was restored by ectopic expression of MltC, while the growth defect at pH 4.7 was not complemented (Fig. 5B). These results indicate that MltC is also essential for bacterial growth under acidic conditions.

**FIG 5.**
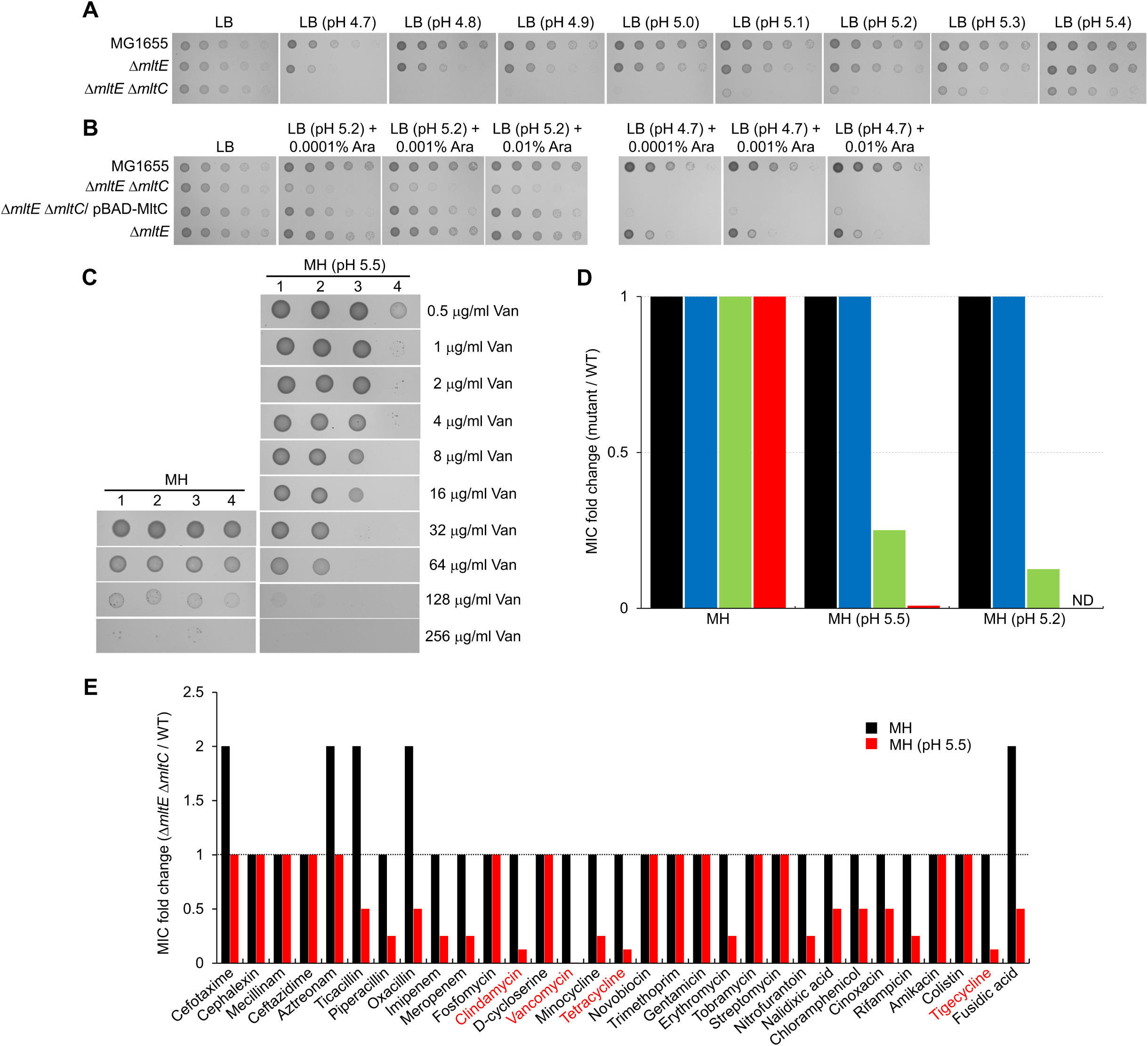
Effects of MltC on bacterial growth and antibiotic resistance at acidic pH. (A) Significantly increased sensitivity of the *mltE* mutant to acidic pH upon additional deletion of the *mltC* gene. The wild-type and indicated mutant cells were serially diluted from 10^8^ to 10^4^ cells/ml in 10-fold steps and spotted onto an LB plate or acidified LB plates at the indicated pHs. (B) Complementation of acid sensitivity of the *mltE mltC* double mutant. Cells of the indicated strains were serially diluted from 10^8^ to 10^4^ cells/ml in 10-fold steps and spotted onto an LB plate and LB plates at the indicated pHs containing the indicated concentrations of arabinose (Ara). (C) Significantly increased sensitivity of the *mltE* mutant to vancomycin upon additional deletion of the *mltC* gene. The MICs of vancomycin (Van) against MG1655 (1), Δ*mltC* (2), Δ*mltE* (3), Δ*mltE* Δ*mltC* (4) strains were measured in Mueller-Hinton (MH) broth plates and acidified MH plates at pH 5.5 containing vancomycin of final concentrations of 512 μg/ml to 3.9 ng/ml in two-fold serial dilutions, according to the Clinical Laboratory Standards Institute guidelines. (D) Effect of MltC on the MICs of vancomycin at acidic pHs. The MICs of vancomycin were measured against MG1655 (black bars), Δ*mltC* (blue bars), Δ*mltE* (green bars), Δ*mltE* Δ*mltC* (red bars) strains in MH medium or MH medium at pH 5.5 or 5.2. The relative MIC values for the mutant cells compared to those for the MG1655 cells are presented. ND, not determined. (E) The MICs of antibiotics against the *mltE mltC* double mutant at acidic pH. The MICs of various antibiotics were measured against the wild-type and *mltE mltC* double mutant strains in MH medium (black bars) or MH medium at pH 5.5 (red bars). The relative MIC values for the *mltE mltC* double mutant cells compared to those for the wild-type cells are presented.

Deletion of the *mltE* gene resulted in increased susceptibility to vancomycin (Fig. 5C), which prompted us to investigate whether additional depletion of MltC in the *mltE* mutant further increases susceptibility to vancomycin. The MIC of vancomycin at pH 5.5 was 4-fold lower in the *mltE* mutant than in the wild-type strain, whereas it was 128-fold lower in the *mltE mltC* double mutant than in the wild-type strain (Fig. 5C and D). These results suggest that MltC also contribute to vancomycin resistance at acidic pH. The high sensitivity of the *mltE mltC* double mutant to vancomycin led us to investigate whether the susceptibilities to other antibiotics are also affected by the deletion of both the genes. In MH broth, there was no antibiotic for which the MIC decreased in the *mltE mltC* double mutant compared to that in the wild-type strain. However, in MH broth at pH 5.5, the MICs of many antibiotics were significantly decreased in the *mltE mltC* double mutant (Fig. 5E). Particularly, the MICs of vancomycin, clindamycin, tetracycline, and tigecycline were more than 8-fold lower in the *mltE mltC* double mutant than in the wild-type strain. Collectively, these experimental data indicate that MltE and MltC are necessary for cell growth and antibiotic resistance at acidic pH.

### MltE and MltC are required for normal cell separation at acidic pH

As several PG hydrolases play an important role in cell morphology (13, 20), we examined the morphology of the *mltE mltC* double mutant at acidic pH. The *mltE mltC* double mutant exhibited normal morphology in LB medium, but notably, the mutant cells displayed a cell separation defect in LB medium at pH 5.5 (Fig. 6). Nuclear staining with DAPI and membrane staining with FM4-64, an amphiphilic membrane dye, clearly revealed separated chromosomes and unseparated septa in the filamentous cells, indicating a chaining phenotype (Fig. 6). Therefore, these results suggest that MltE and MltC are required for daughter cell separation at acidic pH.

**FIG 6.**
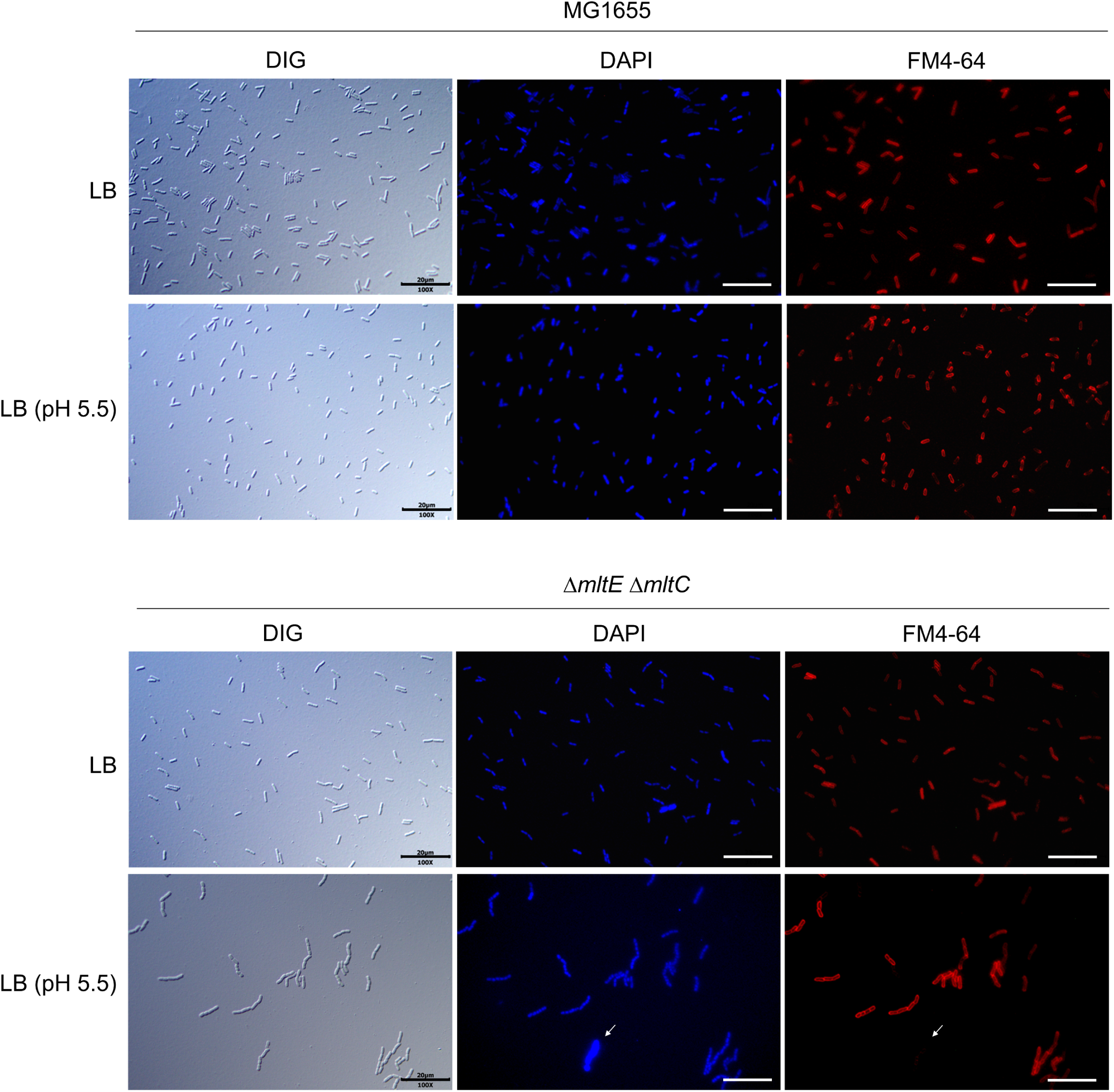
Simultaneous inactivation of MltE and MltC induces filamentous morphology at acidic pH. The wild-type and *mltE mltC* double mutant cells grown in LB medium or acidified LB medium at pH 5.5 to the early exponential phase (OD_600_ = 0.4) were stained with DAPI (blue) and FM4-64 (red), and then spotted on a 1% agarose pad. The arrow indicates a cell with strong DAPI and weak FM4-64 signals. Scale bars, 20 μm.

The chaining phenotype observed in the *mltE mltC* double mutant was reminiscent of mutants defective in the PG amidases, AmiA, AmiB, and AmiC, which are required for daughter cell separation (20, 21). To investigate the relationship between MltE/MltC and PG amidases, we constructed an *mltE mltC* double mutant with additional deletions of PG amidases. Previous studies have shown that in LB medium, the Δ*amiA* mutant exhibits no morphological changes, whereas the Δ*amiA* Δ*amiB* and Δ*amiA* Δ*amiB* Δ*amiC* mutants exhibit a slight and strong chaining phenotypes, respectively (17, 20). These phenotypes were also observed in the Δ*mltE* Δ*mtlC* background in LB medium (Fig. 7). At acidic pH, the chaining phenotype of the Δ*mltE* Δ*mtlC* mutant progressively worsened with additional deletions of PG amidases (Fig. 7). In particular, the Δ*mltE* Δ*mtlC* Δ*amiA* Δ*amiB* mutant exhibited a strong chaining phenotype, only at acidic pH. These results indicate that MltE, MltC, and PG amidases together contribute to daughter cell separation under acidic pH conditions.

**FIG 7.**
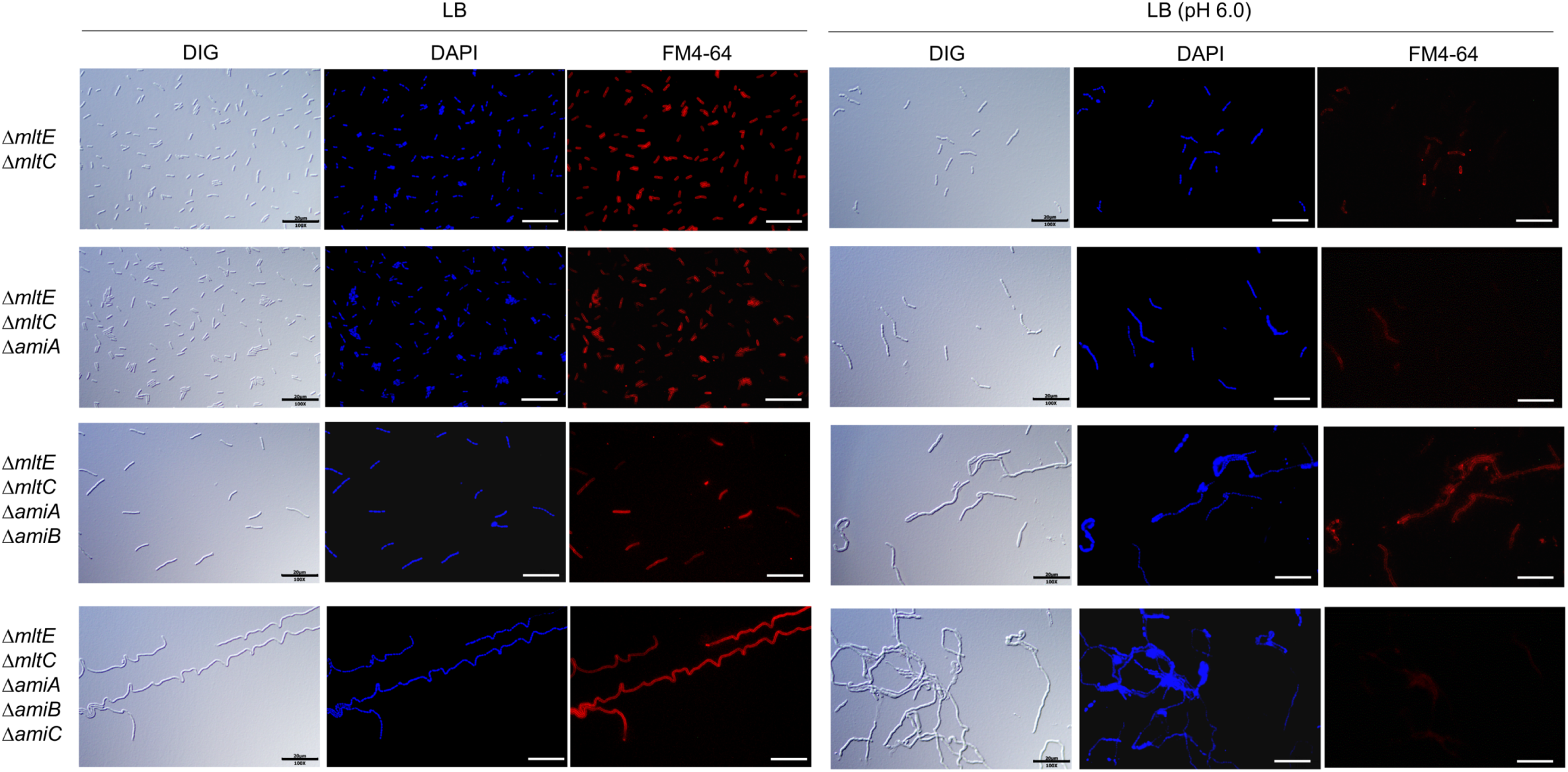
The filamentous phenotype of the *mltE mltC* double mutant at acidic pH is worsened by additional disruptions of PG amidases. The wild-type and indicated mutant cells grown in LB medium or acidified LB medium at pH 6.0 to the early exponential phase (OD_600_ = 0.4) were stained with DAPI (blue) and FM4-64 (red), and then spotted on a 1% agarose pad. Scale bars, 20 μm.

### Simultaneous inactivation of MltE and MltC results in increased membrane permeability at acidic pH

When analyzing the morphological features of the *mltE mltC* double mutant, we observed cells (indicated by arrows) that exhibited strong DAPI signals (Figs. 6 and 8A). Notably, these cells were not stained with FM4-64 dye (Figs. 6 and 8A) (22). At pHs 5.1 and 4.8, nearly all the cells of the *mltE mltC* double mutant displayed strong DAPI and weak FM4-64 signals (Fig. S1). These staining patterns were not observed in the wild-type strain even at pH 4.8 (Fig. S1). These results suggest that the *mltE mltC* double mutant may exhibit increased membrane permeability at acidic pH. To explore this further, we examined the effects of MltE and MltC on envelope stress responses. Membrane permeability was assessed using the cell-impermeable fluorescent DNA dye, SYTOX green (22, 23). At acidic pH, SYTOX green signals were not detected in the wild-type strain, but strong intracellular SYTOX green signals were observed in many cells of the *mltE mltC* double mutant (Fig. 8B). The *mltE mltC* double mutant was also highly sensitive to envelope stresses, including sodium dodecyl sulfate (SDS) and salt stresses, at acidic pH (Fig. 8C). Collectively, these data indicate that the simultaneous inactivation of MltE and MltC induces increased membrane permeability at acidic pH.

**FIG 8.**
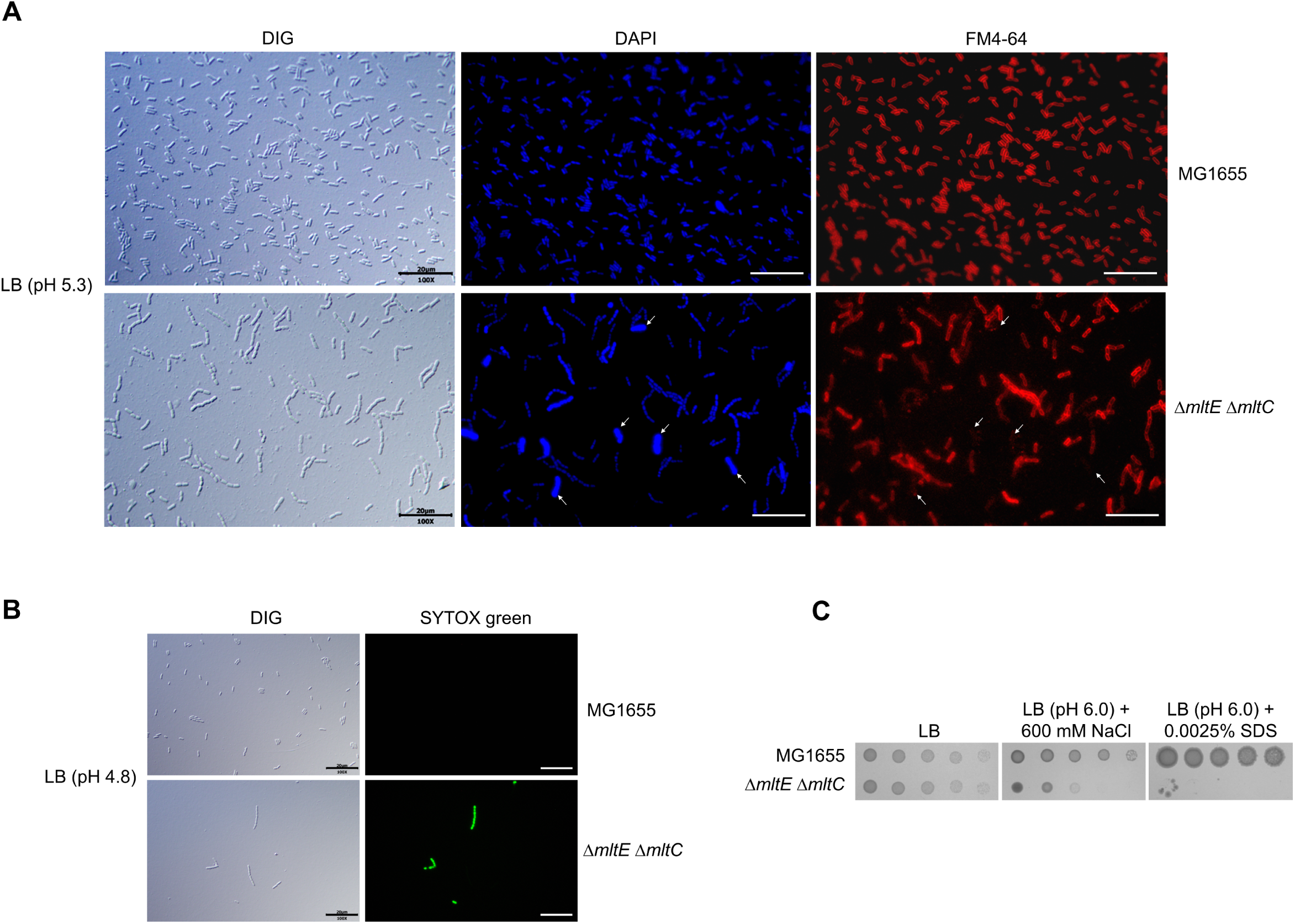
The *mltE mltC* double mutant exhibits increased membrane permeability. (A) Cells of the *mltE mltC* double mutant with strong DAPI and weak FM4-64 signals at acidic pH. The wild-type and *mltE mltC* double mutant cells grown in LB medium at pH 5.3 to the early exponential phase (OD_600_ = 0.4) were stained with DAPI (blue) and FM4-64 (red), and then spotted on a 1% agarose pad. The arrows indicate cells with strong DAPI and weak FM4-64 signals. Scale bars, 20 μm. (B) Intracellular penetration of a SYTOX green dye in the *mltE mltC* double mutant at acidic pH. The wild-type and *mltE mltC* double mutant cells grown in LB medium at pH 4.8 to the early exponential phase (OD_600_ = 0.4) were stained with SYTOX green (green), and then spotted on a 1% agarose pad. Scale bars, 20 μm. (C) Salt and SDS sensitivities of the *mltE mltC* double mutant at acidic pH. The cells of the wild-type and *mltE mltC* double mutant strains were serially diluted from 10^8^ to 10^4^ cells/ml in 10-fold steps and spotted onto an LB plate and an LB plate at pH 6.0 containing 600 mM NaCl or 0.0025% SDS.

### Inhibition of LTG activities increases the antimicrobial activity of vancomycin only at acidic pH

Antibiotic adjuvants that promote the antimicrobial activity of conventional antibiotics have the potential to overcome antibiotic resistance of Gram-negative bacteria. We turned to a previously developed small molecule with potent LTG inhibitory activity and tested it in combination with vancomycin. The phenothiazinium dye thionine acetate, a weak inhibitor of LTGs (24), exhibited a potentiator effect of vancomycin only at acidic pH (Fig. S2). In MH broth, the presence of thionine acetate did not change the MIC of vancomycin, but at pH 5.2, the MIC of vancomycin was 16-fold lower in the presence of thionine acetate than in the absence of thionine acetate. As a control, we tested the effect of thionine acetate on fosfomycin susceptibility. The MIC of fosfomycin was two-fold lower in the presence of thionine acetate than in the absence of thionine acetate regardless of medium pH (Fig. S2). Therefore, these results showed the potential significance of LTGs as important targets for the development of potentiators to augment the antimicrobial efficacy of antibiotics in acidic pH environments.

## DISCUSSION

The periplasm of Gram-negative bacteria constitutes a challenging intracellular space that is directly impacted by extracellular stress conditions, such as acidic stress, without effective homeostatic regulation. A limited number of enzymes operate within the periplasm, with PG hydrolases serving as the prime examples. In this study, we elucidated that a diverse array of LTGs contribute to maintaining their functions in the periplasm under acidic stress. Our findings underscore the importance of MltE and MltC, representative LTGs, in bacterial growth, cytokinesis, and antibiotic resistance under acidic stress, highlighting the physiological significance of LTG redundancy.

The pronounced redundancy of PG synthases and hydrolases functioning in the periplasm allows bacteria to overcome challenging periplasmic stress (11). Notably, several acidic enzymes essential for overcoming acidic stress have been identified (1, 11, 12). Among the PG DD-carboxypeptidases, DacD is an acidic enzyme with significantly enhanced enzymatic activity that is crucial for maintaining morphology under acidic stress (11). ActS-mediated stimulation of PG amidase activity at acidic pH contributes to cell separation under acidic stress (12). The class A penicillin-binding protein PBP1b is also an acidic pH specialist involved in PG synthesis. It’s uninhibited glycosyltransferase and transpeptidase activities at low pH essentially regulate bacterial growth under these conditions (1). In this study, we revealed that MltE and MltC, representatives of LTGs, are also acidic pH specialists and play vital roles in cell growth and daughter cell separation at acidic pH. These findings underscore the presence of periplasmic enzymes in bacteria that are specially adapted to overcome acidic stress.

Recent studies highlighted significant changes in the protein levels of PG-degrading enzymes in response to extracellular conditions (11, 13, 25). For instance, the protein levels of DacD and DacC, members of the PG DD-carboxypeptidase family, are notably enhanced at acidic and alkaline pH conditions, respectively (11, 13). Among PG endopeptidases, the protein levels of MepS and MepM are regulated by the availability of amino acids, particularly aromatic amino acids (25). Our data demonstrated that among LTGs, MltC levels increased under acidic pH conditions (Fig. 4). These findings suggest that protein levels of other PG-degrading enzymes may also be regulated under harsh periplasmic stress conditions.

PG-degrading enzymes are strongly linked to antibiotic resistance, particularly against PG-targeted antibiotics such as β-lactams and vancomycin. PG DD-carboxypeptidases, especially DacA, exhibit divergent effects on β-lactams and vancomycin (26–28). Depletion of DacA renders *E. coli* sensitive to most β-lactams and induces strong resistance to vancomycin (26). In the DacA-defective mutant, vancomycin resistance is attributed to an increased abundance of decoy D-alanine-D-alanine residues within PG, but the exact mechanism of β-lactam sensitivity remains unknown (26). PG LD-transpeptidases are closely associated with β-lactam resistance in various pathogenic bacteria, including *Mycobacterium tuberculosis* and *Clostridium difficile* (29). These bacteria predominantly feature PG with 3-3 cross-links formed by L,D-transpeptidases, making them resistant to β-lactams that inhibit DD-transpeptidases but not LD-transpeptidases. Several LTG enzymes have also been implicated in β-lactam resistance. Depletion of Slt induces β-lactam sensitivity in *E. coli* (8, 30). In *Pseudomonas aeruginosa*, strains lacking multiple LTGs, such as Slt, exhibit increased sensitivity to β-lactams (31). Our results demonstrated that the loss of both MltE and MltC increased the sensitivity to many antibiotics under acidic conditions (Fig. 5E). Depletion of these proteins resulted in a strong sensitivity to vancomycin, clindamycin, tetracycline, and tigecycline. These phenotypes appeared to be partially caused by increased membrane permeability (Fig. 8). Collectively, our findings suggest that stress-specific PG-degrading enzymes play significant roles in antibiotic resistance under specific environmental pressures.

Cytokinesis in bacterial cells requires both PG synthesis via the divisome complex (32) and PG degradation via PG amidases (20), highlighting their crucial roles in this process. Previous studies have shown that LTGs also contribute to bacterial cytokinesis (6, 7). In this study, we demonstrated that MltE and MltC are involved in bacterial cytokinesis under acidic conditions (Fig. 6). Importantly, their roles in cytokinesis were independent of the functions of PG amidases (Fig. 7). These results suggest that LTGs play a distinct role in cytokinesis that cannot be substituted by PG amidases.

Our findings suggest that MltE and MltC are promising targets for developing potentiators that enhance the antimicrobial activities of conventional antibiotics under acidic pH conditions. Bacteria encounter acidic pH within the host, such as in phagolysosomes of phagocytes and during passage through the stomach or small intestine (33). Consequently, inhibition of MltE and MltC could augment the elimination of bacteria within phagolysosomes when combined with conventional antibiotics. This strategy has the potential to preserve or improve the therapeutic efficacy of clinically relevant antibiotics.

## MATERIALS AND METHODS

### Bacterial strains, plasmids, and culture conditions

All bacterial strains and plasmids used in this study are listed in Table S1. *E. coli* strains were cultured in Luria-Bertani (LB) medium (pH 7.4) at 37°C, unless otherwise stated. Antibiotics at 100 μg/ml (ampicillin), 50 μg/ml (kanamycin), 5 μg/ml (chloramphenicol), or 10 μg/ml (tetracycline) concentration were added to the culture medium when necessary. To measure bacterial growth, cells that were cultured overnight in LB medium were inoculated into fresh LB medium. When the OD_600_ reached approximately 0.8, the cultured cells were diluted from 10^8^ to 10^4^ cells/ml in 10-fold dilutions and 2 μl of aliquots were spotted onto LB plates or LB plates containing the indicated reagents. After 10–20 h of incubation at 37°C, the plates were imaged using the digital camera, EOS 100D (Canon Inc., Japan). LB plates with acidic pHs were made by adding sodium citrate (50 mM, final concentration) to the LB medium and adjusting the pH to acidic values using 10 N HCl solution.

LT-depleted strains were constructed by homologous recombination with λ red recombinase, as previously described (34). Deletion cassettes were prepared by polymerase chain reaction (PCR) using primers with a 50 bp sequence for homologous recombination (See Table S2) and the pKD13 plasmid as a template. After PCR purification, the PCR products were inserted into MG1655 cells harboring pKD46 plasmids expressing λ red recombinase. LT-depleted cells were selected on LB plates containing kanamycin. Deletion of the target gene was confirmed by PCR using confirming primers (Primer sequences in Table S2). Finally, the kanamycin resistance gene was removed by FLP-FRT recombination (34). The pCP20 plasmid expressing FLP recombinase was inserted into LT-depleted cells, and the removal of the kanamycin resistance gene was confirmed by PCR using the confirming primers mentioned above. The pCP20 plasmid curing was performed at 37°C, and not 42°C, to minimize the physiological effect on the deletion strain, as previously reported (13, 14).

The pBAD24 or pET28a plasmid expressing LTGs, such as MltE, was constructed using infusion cloning (Clontech, USA), as previously reported (35). The sequence of the entire open reading frame of the target gene was amplified by PCR using cloning primers (Primer sequences in Table S2) and genomic DNA of MG1655 as a template. After PCR purification, the PCR product was inserted into the pBAD24 plasmid digested with XbaI and EcoRI or the pBAD24 plasmid digested with NdeI and BamHI. The PCR product was inserted into the plasmid by homologous recombination between overlapping sequences using infusion cloning. The exact insertion of the target gene was confirmed using DNA sequencing. The pBAD-MltE(E64Q) plasmid was constructed by PCR using point-mutant primers and the pBAD-MltE plasmid as a template. After PCR purification, the original plasmid template was digested overnight with the DpnI restriction enzyme. The point mutation in the *mltE* gene was confirmed by DNA sequencing. The pBAD-MltE(S17D&S18D)-Flag plasmid was constructed by the same method. To construct the pBAD24 plasmid expressing chimeric MltE proteins with the signal sequence of DsbA (amino acids 1–19) and a Flag-tag, the *mltE* gene was cloned into the pBAD-DsbA(ss)-Flag plasmid, resulting in the pBAD-DsbA(ss)- MltE-Flag plasmid.

To measure intracellular protein levels of LTGs, a *3×Flag* gene was inserted into the chromosomal 3’ region of the target gene using λ red recombinase. DNA cassettes for homologous recombination were prepared by PCR using the primer sets listed in Table S2 and the pBAD-Flag-FRT-Kan plasmid, which contains both the 3×Flag gene and the FRT-flanked kanamycin resistance gene, as a template. After the removal of template plasmids through overnight DpnI treatment, the PCR products were inserted into the MG1655 strain harboring the pKD46 plasmid. The insertion of both the 3×Flag gene and kanamycin resistance gene into the chromosomal 3’ region of the target gene was confirmed by PCR. Kanamycin resistance genes were removed as described above.

### Quantitative real-time PCR

MG1655 cells were cultured in LB medium or acidified LB medium that had a pH 4.7. When the OD_600_ reached approximately 0.4, total RNA of cells was extracted using an RNeasy Mini Kit (Qiagen, USA). Samples were then incubated with RNase-free DNase I (Promega, USA) at 37°C for 2 h, to remove genomic DNA. All RNA in the extract samples were converted into cDNA using the cDNA EcoDry Premix (Clontech, USA). Quantitative real-time PCR was performed using a CFX96 Real-Time System (Bio-Rad, USA). PCR was performed in SYBR Premix Ex Taq II (Takara, Japan) solution using MltE RT-PCR primers (See Table S2) and 10-fold diluted cDNA samples as templates. The expression level of *mltE* was calculated based on the 16S rRNA reference gene.

### Detection of intracellular proteins levels of LTGs

The protein levels of LTGs (MltE, MltG, and MltC) were measured using MG1655 strains with chromosomal insertion of the *3×Flag* gene into the 3’ region of each gene (Table S1). Each strain was cultured in LB medium or acidified LB medium with pH 4.7 at 37°C. When the OD_600_ reached approximately 0.4, cells were harvested and SDS sample buffer was added. After boiling for 5 min at 100°C, total cell proteins were separated on 4–20% Tris–glycine polyacrylamide gels (KOMA Biotech, Korea). After transferring all proteins onto polyvinylidene fluoride (PVDF) membranes, the protein levels of LTGs and DnaK were determined using monoclonal antibodies against Flag-tag (Santa Cruz Biotechnology, USA) and anti-DnaK (Abcam, USA), respectively, according to standard western blot procedures. The protein level of DnaK was used as loading control.

### Purification of LTG proteins

ER2566 cells harboring the pET28a-based plasmids expressing His-tagged LTGs were cultured in LB medium at 37°C. At an OD_600_ of approximately 0.5, 1 mM (final concentration) isopropyl-β-D-1-thiogalactopyranoside (IPTG) was added into culture medium and cells were cultured overnight at 16°C. Harvested cells were resuspended in 3 ml of binding buffer (50 mM Tris-HCl [pH 8.0] and 200 mM NaCl) and disrupted using a French press at 10,000 psi. Soluble proteins of cell extracts were separated by centrifugation at 10,000 × *g* for 20 min at 4°C. The supernatants from the samples were added to 0.5 ml of a Talon metal affinity resin (Clontech, USA) equilibrated with binding buffer. After washing three times with the binding buffer, His-tagged LTG proteins bound to the resin were eluted using a binding buffer containing 150 mM imidazole. To remove the imidazole, the eluted samples were dialyzed overnight in 2 L of 50 mM Tris-HCl buffer (pH 8.0) containing 50 mM NaCl. The dialyzed samples were promptly subjected to measurement of the enzymatic activity of the LTGs, without freezing.

### Purification of PG

Wild-type cells were cultured in 1 L of LB medium until the early exponential phase and harvested. After resuspension in 20 ml of phosphate buffered saline (PBS), 80 ml of 5% SDS solution was added, and the mixture was boiled at 100°C for 30 min. After overnight incubation at room temperature, the mixture was ultracentrifuged using a Himac Micro Ultracentrifuge CS120FNX (Hitachi, Japan) at 165,000 × *g* for 1 h at room temperature. After removing the supernatant, the pellet was washed six times with distilled water (DW) to remove SDS. α-Chymotrypsin (final concentration of 300 μg/ml) and α-amylase (final concentration of 0.2 mg/ml) were added to the pellet sample resuspended in 1 ml of PBP buffer, to remove proteins and glycogen. After overnight incubation at 37°C, SDS (final concentration of 1%) was added and then the sample was boiled at 100°C for 30 min. After ultracentrifugation at 260,000 × *g* for 20 min at room temperature, the pellet was washed with DW at least four times to remove the remaining SDS. Purified PG was resuspended in 1 ml of DW and was stored at −80°C until further experiments.

### Assessment of the enzymatic activity of LTGs

Purified PG (1 ml) was labeled by mixing with RBB (20 mM, final concentration) in 0.25 N NaOH and incubating the mixture at 37°C for 6 h. After neutralization with 10 mM HCl (final concentration), the samples were centrifuged at 21,000 × *g* for 20 min at room temperature. The pellet was washed repeatedly with PBS until the supernatant cleared and resuspended in 10 ml of PBS. To assess the enzymatic activity of the purified LTG proteins, RBB-labeled PG (100 μl) was mixed with 500 μg/ml of purified LTGs in 50 mM potassium phosphate (pH 7.0) or 50 mM potassium citrate (pH 5.5 or 5.0) buffer to a final volume of 700 μl. After incubation at 37°C for 2 h, the enzymatic reaction was terminated by adding 300 μl of ethanol. After centrifugation at 21,000 × *g* for 20 min at room temperature, the absorbance of the supernatant was measured at 595 nm.

### Determination of MIC values

The MIC values of antibiotics were determined in Mueller-Hinton (MH) broth or acidified MH broth at pH 5.5 or 5.2, according to the Clinical Laboratory Standards Institute guidelines (36). The cells grown in LB medium overnight were cultured in MH broth to a McFarland turbidity standard of 0.5 (approximately 1.5 × 10^8^ cells/ml). After dilution with MH broth to a final concentration of 10^7^ cells/ml, 10 μl of the diluted samples was spotted onto MH plates with the indicated pH values and containing antibiotics at final concentrations ranging from 1024 μg/ml to 7.8 ng/ml in two-fold serial dilutions. MH plates with acidic pH were made by adding sodium citrate (50 mM, final concentration) to the MH medium and adjusting the pH to acidic values using 10 N HCl solution. After incubation at 37°C for 20 h, the MIC value of each antibiotic, which is the lowest concentration of antibiotic that inhibits a lawn growth of cells, was determined.

### Evaluation of cell shape using microscopy

The cells were cultured in LB medium or an acidified one of the indicated pH, to an OD_600_ of approximately 0.4. The cells were stained with 5 μg/ml FM4-64 [N-(3-triethylammoniumpropyl)-4-(*p*-diethylaminophenylhexatrienyl)-pyridinium dibromide] and 5 μg/ml DAPI (4’,6-diamidino-2-phenylindole) for 10 min at room temperature. The stained cells were spotted onto a 1% agarose pad with PBS buffer and the cells on the pad were observed using an Eclipse Ni microscope (Nikon, Japan).

### Determination of membrane integrity using fluorescent staining

Cells of the wild-type and *mltE mltC* double mutant strains were cultured in LB medium at the indicated pH to an OD_600_ of approximately 0.1. Cells from 1 ml bacterial suspension were harvested and resuspended in 100 μl of LB medium. The resuspended cells were stained with 5 μg/ml FM4-64 and 5 μg/ml DAPI or 0.5 μM Sytox green and were spotted onto a 1% agarose pad with PBS buffer. The cells on the pad were observed under an Eclipse Ni microscope.

